# Diagnosed and undiagnosed Diabetes mellitus among urban adults: a population based cross-sectional study

**DOI:** 10.1101/532705

**Authors:** Behailu Hawulte Ayele, Hirbo Shore, Addisu Shunu, Melkamu Merid Mengesha

## Abstract

**Background:** Globally, diabetes mellitus (DM) accounts for 8.8% (424.9 million) morbidity and 4 million deaths. In 2017, more than 79% of people with diabetes live in low- and middle- income countries. To this end, locally available evidence can identify target groups for intervention. However, in resource-poor settings, population-based evidence on diabetes prevalence and on its risk factors is lacking. This study, therefore, assessed prevalence of Diabetes mellitus and associated factors among adults living in Dire Dawa town, Eastern Ethiopia.

**Methods:** A total of 782 data points were analyzed from a random sample of the adult population aged 25-64 years who lived in Dire Dawa. World health organization STEPwise approach to non-communicable disease risk factors surveillance (WHO NCD STEPS) instrument was used to collect data. We estimated undiagnosed DM, uncontrolled DM among existing cases and the overall prevalence of DM. Hierarchical logistic regression models were run to identify correlates of diabetes mellitus, and STATA v 14.2 was used for data management and analysis. All statistical tests were declared significant at p-value<0.05.

**Results:** The prevalence of DM among adults aged 25-64 was 8.95% (95% confidence interval (CI): 7.1, 11.2) and the magnitude of undiagnosed DM was 3.3% (95% CI: 2.3, 4.8). The magnitude of uncontrolled DM among those taking DM medications during the survey was 1.4% (95% CI: 0.8, 2.5). The prevalence of DM was 2.3 times more likely among the age group of 55-64 years (Adjusted Odds Ratio (AOR) 95% CI: 1.1, 5.0). Similarly, consuming two or less serving of vegetables/week increased the risk of DM, (AOR=2.1, 95% CI: 1.1, 2.9). Maintaining normal body mass index level was negatively correlated with the risk of DM, (AOR=0.6, 95% CI: 0.3, 0.8).

**Conclusion:** The overall prevalence of diabetes mellitus was relatively high, and the magnitude of undiagnosed DM was a great concern. Therefore, creating community awareness, regular blood sugar checking, appropriate weight control and, increased consumption of vegetables would be helpful in preventing incident cases of DM.

## Introduction

Diabetes mellitus is a chronic disease caused by an inherited and/or acquired deficiency in the production of insulin by the pancreas, or by the ineffectiveness of the insulin produced [1, 2]. The global prevalence of diabetes among adults over 18 years of age has risen from 4.7% in 1980 to 8.8% in 2017 (3-5). In 2015, an estimated 4.0 million deaths were directly caused by diabetes [3]. Almost half of all deaths attributable to high blood glucose occur before the age of 70. World Health Organization (WHO) projected that diabetes will be the seventh leading cause of death by 2030. In 2045, Diabetes is projected to affect 628.6 million people worldwide [2, 3, 6].

Diabetes prevalence has been rising more rapidly in low- and middle-income countries. According to the IDF report of 2017, more than 79% of people with diabetes live in low- and middle- income countries [3, 7]. The African region (according to IDF regional divisions) has the lowest prevalence in 2017, likely due to lower levels of urbanization, under-nutrition, lowers levels of obesity and higher rates of communicable diseases. However, the proportion of all deaths due to diabetes occurring before age 60 amounts to around 77.0% in the region [8].

In Ethiopia, a country in Sub-Saharan Africa, the magnitude of diabetes is alarming. According to world health organization diabetes country profiles of 2016, 3.8% (4.0% among male and 3.6% among females) of the population had diabetes morbidity. In the country, diabetes accounts for 1% of overall mortality [4]. The national report of 2016 STEPS survey showed, 5.9% (6.0% in male & 5.8% in females) of Ethiopians had raised blood glucose level and 98.4% had at least on risk factors[9].

An estimated gross domestic product loss due to diabetes, including both the direct and indirect costs will total US$ 1.7 trillion, where LMICs will share a total of US$ 800 billion owing to the unacceptably high burden of diabetes. Besides the economic burden on the health-care system and national economy, diabetes imposes catastrophic out-of-pocket personal expenditures from loss of income due to disability and premature death [10]. Diabetes is also a major cause of blindness, kidney failure, heart attacks, stroke, amputation and, death. The negative impacts of diabetes are unacceptably high as there are effective measures of public health and clinical preventions including a healthy diet, regular physical activity, maintaining normal body weight and avoiding tobacco use (1).

Prior community-based studies across the Globe reported the prevalence of diabetes mellitus ranges from 1.4 to 19.1% [9, 11–29]. Factors associated with the occurrence of diabetes included: Being older age [11–15, 18, 19, 21–23, 25, 28], lower education level [14, 21], being unemployed [13], living in an urban setting [9], having a family history of DM [12, 18, 22, 25], low fruit and vegetable consumption[30–33], lower levels of physical activity[12, 14, 18, 21, 23, 27, 28], smoking [12, 14, 23, 29], alcohol consumption [9, 16, 23], khat chewing[9], being overweight[17, 19, 22, 23, 27, 29], obesity [11, 13–17, 19, 21–24, 27, 29], Waist to hip ratio (WHR)[28, 29], dyslipidemia [13, 14, 25, 29] and being hypertensive [13, 14, 17, 19, 22, 25, 28, 29] are some of the factors that were associated with the occurrence of Diabetic mellitus.

Previously, even though a number of studies were conducted on the prevalence of Diabetes mellitus and its risk factors, there are few studies conducted at the population level in Ethiopia. To our knowledge, population-level research has not been conducted in the current study area. In addition, previous research was done in different parts of the world [10–14, 17, 18, 20–22, 24, 27] on the prevalence of diabetes mellitus analyzed independent variables at an equal level as a risk factor without considering the multidimensional nature of disease risk factors. Considering these gaps, the current study assessed the prevalence of Diabetes mellitus and its multi-level risk factors.

## Methods and Materials

### Study setting and design

A community-based cross-sectional study was conducted from June 01-21, 2017 in Dire Dawa City Administration. Dire Dawa city is located in central eastern Ethiopia, 30 miles (48 km) northwest of Harar. It lies at the intersection of roads from Addis Ababa, and Djibouti. It is located 515 kilometers east of Addis Ababa, the capital of Ethiopia. The Dire Dawa administrative council consists of the city of Dire Dawa and the surrounding rural areas. The council has 4 ‘Keftegnas’, 9 urban kebeles and 28 rural peasants associations. According to the Central Statistical Authority report of 2013, the total population of the administration was 405,444 with 4,530/km^2^ (11,700/sq mi) population density, among the total population, 196,777 were male and 208,666 and female [34].

### Populations

This study used the sample calculated for a study on metabolic syndrome in the study setting (unpublished). The total sample was 903, and only 782 valid observations were maintained for this analysis. The study was conducted among adult populations aged 25-64 years who had lived in Dire Dawa for at least six months. Multi-stage sampling technique was used with the primary sampling unit of being 5 kebeles (the lowest administrative unit, equivalent to sub-district) from the total 9 kebeles. The sample size was proportionally distributed to each of kebeles based on the number of the household. Finally, a systematic random sampling technique was employed to select households to be visited. During the visit, the number of adults aged between 25 and 64 was identified, and if two or more eligible adults were found in a household, the lottery method was used to select one adult for the interview.

### Instruments and Data collection

This study was used the World health organization STEPwise approach to non-communicable disease risk factors surveillance (WHO NCD STEPS) instrument which consisted of three steps for measuring the risk of the non-communicable diseases risk factors. The first step consists of a collection of core and expanded socio-demographic and behavioral characteristics of the study population. The second step involves core and expanded physical measurement and the third step consists of biochemical measurement[35]. The questionnaire was translated to Amharic and Afan Oromo, the two widely spoken languages in Dire Dawa. Data were collected by health care professionals who had a Bachelor of Science degree (BSc) in nursing and intensive training on the study objectives, the overall STEP survey procedure and, study tool was given for data collectors. A pretest was conducted in the kebele that was not included in the study to check for the validity of the instruments and necessary adjustments were made before the actual data collection.

The anthropometric measurement was carried out following standard procedures and using calibrated instruments[36]. Weight was measured using a standard digital scale. We used stadiometer to measure height and results were recorded to the nearest 0.5 cm. Digital BP apparatus was used to measure blood pressure. The blood pressure was measured three times 3 to 5 minute apart from the left arm while the subject is in sitting position. The averages of the last two measures were used to determine blood pressure. Digital glucometer was used to measure capillary blood sugar after participants were asked the time lapsed from the last meal.

### Operational definition

Diabetes Mellitus: according to WHO and International Diabetic Association (IDA) diabetes mellitus was diagnosed if one or more of the following criteria are met: Fasting plasma glucose ≥126 mg/dL or random glucose > 200 mg/dL(8). Subjects who were taking DM medications during the survey were also identified as previously diagnosed with DM.

Hypertension: defined as persistent systolic blood pressure≥140mmHg or diastolic blood pressure ≥90mmHg or reported use of anti-hypertensive medication [35, 37–39].

BMI: is defined as a person’s weight in kilograms divided by the square of the person’s height in meters (kg/m2) and categorized as follow: BMI < 18.5; underweight, BMI 18.5-24.9; normal and BMI ≥25; overweight and obese [40].

### Data processing and Analysis

The data were entered into Epi-Data version 3.1 and exported to STATA v 14.2 statistical software. The hierarchical nature of chronic diseases risks factors were considered in the analysis to identify factors associated with raised blood glucose at a different level of risk, from the most distal to the proximal causal risk factors[35, 41]. In line with this, three binary logistic regression models were run to identify factors associated with DM considering risk factors operating at a different level. At each level, variables with a p-value less than 0.3 were considered for multivariate binary logistic regression. The first model was adjusted only for socio-demographic characteristics (such as sex, age, educational status, marital status, and ethnicity). The second model was adjusted for sociodemographic characteristics and behavioral characteristics (such as consumption of fruits and vegetable, and exercises). The last model adjusted for risk factors in socio-demographic characteristics, behavioral characteristics and immediate factors (such as hypertension and BMI)[42]. An odds ratio with a 95% confidence interval was presented and fitness-test was checked for each model.

Multiple imputations by chained equations (MICE) were used to do imputation assuming missing at random. Before imputation, we dropped 32 observations for which blood glucose, the outcome variable for this analysis, was missing. Similarly, 65 observations were deleted for a total of 21 variables which had less than at most 7 missing values. Models used for imputing missing values of variables include binary logistic regression (type of blood glucose measure, fasting versus random, alcohol; consumption within the past 30 days), multinomial logistic regression (ethnicity), predictive mean matching (age, number of days eat fruit, servings of fruit per day, number of days eat vegetables, servings of vegetables per day, frequency of moderate intensity activity at work, duration of moderate intensity activity at work, number of days walk or bicycle for at least 10 minutes, duration spent walking or bicycling on a typical day), and linear regression (years in school, height, weight, waist circumference, and hip circumference). Values imputed range from 6% to 41% of the analysis sample, 782. Research evidence indicate that multiple imputation, when data size is adequate, performs well with high level of missing’s, up 50% missing values[43].

## Results

### Socio-demographic characteristics of the study participants

Of the total 903 sample calculated for the metabolic syndrome study, 879 completed the survey. There were several variables with missing values, and we did imputation to prevent loss of study. Before imputation, however, 97 observations of several variables which had missing values of 7 or less values were deleted, resulting in a final analysis sample of 782. The Majority, 66.7% of participants was female and the mean age ± SD for the studied participants was 40.2 + 13.0 years. Five hundred twenty (66.5%) participants were married and 87% had achieved some level of formal education (Table 1).

**Table 1.** Socio-demographic characteristics of adults age 25-65 in Diredawa City, 2017. (n=782).

| Variables |  | Frequency (n=782) | Percentage |
| --- | --- | --- | --- |
| **Sex** | Male | 260 | 33.3 |
|  | Female | 522 | 66.7 |
| **Age** | 25-29 | 214 | 27.4 |
|  | 30-44 | 291 | 37.2 |
|  | 45-54 | 107 | 13.7 |
|  | 55-64 | 170 | 21.7 |
| **Educational status** | No formal education | 102 | 13.0 |
|  | Primary (1-8) | 304 | 38.9 |
|  | Secondary(9-12) | 287 | 36.7 |
|  | College/ University | 89 | 11.4 |
| **Marital status** | Married | 520 | 66.5 |
|  | Not married | 156 | 19.9 |
|  | Divorced | 66 | 8.5 |
|  | Widowed | 40 | 5.1 |
| **Occupational status** | Government employee | 99 | 12.7 |
|  | NGO Employee | 106 | 13.5 |
|  | Merchant | 104 | 13.3 |
|  | Unemployed | 340 | 43.5 |
|  | Other<sup>§</sup> | 133 | 17.0 |
| **Ethnicity** | Amhara | 386 | 49.3 |
|  | Oromo | 268 | 34.3 |
|  | Others* | 128 | 16.4 |

§=Students, Farmers, Housewife's, Pensioners, daily laborer, driver, janitor, and private employee,
\*= Gurage, Somali, Tigraywa, Harari, wolaita, sidama, Djiboutian

### Behavioral, Diet, and physical measurement of the study participants

From the total participants, 91.4% had never smoked cigarettes in their lifetime. Around 30.6% of the participants had drunk alcohol at least once in their lifetime and 24.0% had drunk alcohol in the past 30 days prior to the study. Three hundred thirty-one (42.3%) and 300(38.4%) ate 5 or more serving of fruits and vegetable a week respectively. Concerning physical activities, 324(41.4%) of the participants did not achieve 600-metabolic equivalent tasks (MET-minutes) and the mean ± SD of minutes spent on sitting or reclining per week was 332.9 ± 187.2. The mean ± SD of BMI in the current study was 24.2 ± 4.6 Kg/m^2^ and around 37% of study participants were overweight or obese (Table 2).

**Table 1:**
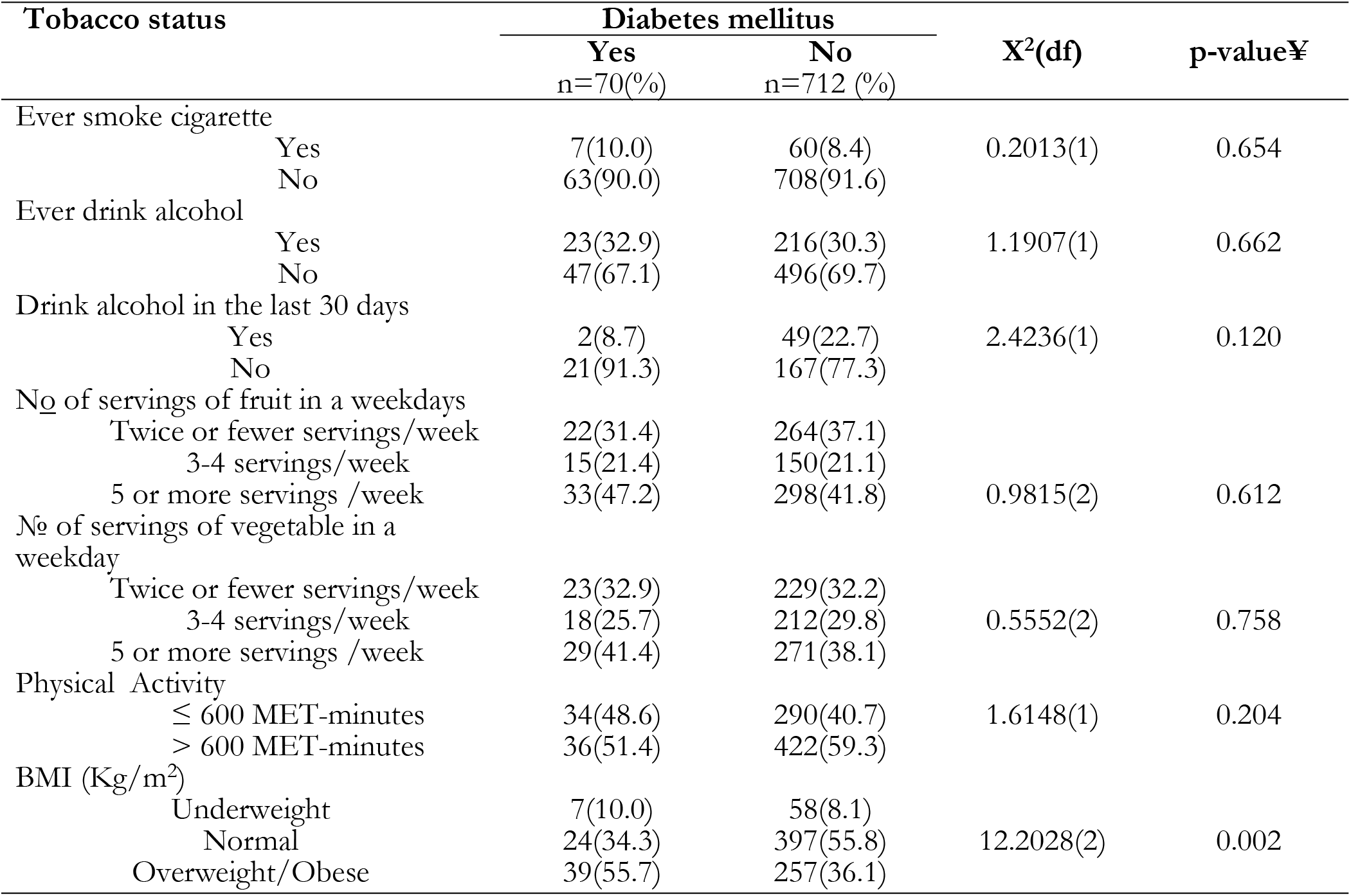
Behavioral, Diet, and physical characteristics of adults age 25-65 in Dire Dawa City, 2017. (n=782)

### The magnitude of diabetes mellitus

The overall prevalence of Diabetes mellitus (DM) among the adult population in Dire Dawa city was 8.95% (95% CI: 7.1, 11.2) and the magnitude of undiagnosed DM was 3.3% (95% CI: 2.4, 4.8). The magnitude of uncontrolled DM among those taking DM medications during the survey was 1.4% (95% CI: 0.8, 2.5). The mean + Standard Deviation (SD) of random blood sugar (RBS) and fasting blood sugar (FBS) measure was 120.7 ± 49.4 and 107 ±22.8 respectively. More than half of the participants, 52.9% had never checked their blood glucose. Among those who ever had checked their blood glucose level, 36 (9.8%) have been told to have raised blood glucose, but only 5.6% (95% CI: 4.2, 7.5) were taking DM medications on the date of the survey.

The prevalence DM was higher among female than male; 9.6% (95% CI: 7.3, 12.4) among female and 7.7% (95% CI: 5.0, 11.6) among male, but the difference was statistically insignificant (chi2 (df) =0.76(1), P-value=0.38). One third (34.3%) of DM cases were between the age of 30-44 years old and 55.7% of DM cases were overweight or obese (Fig. 1).

**Fig.1:**
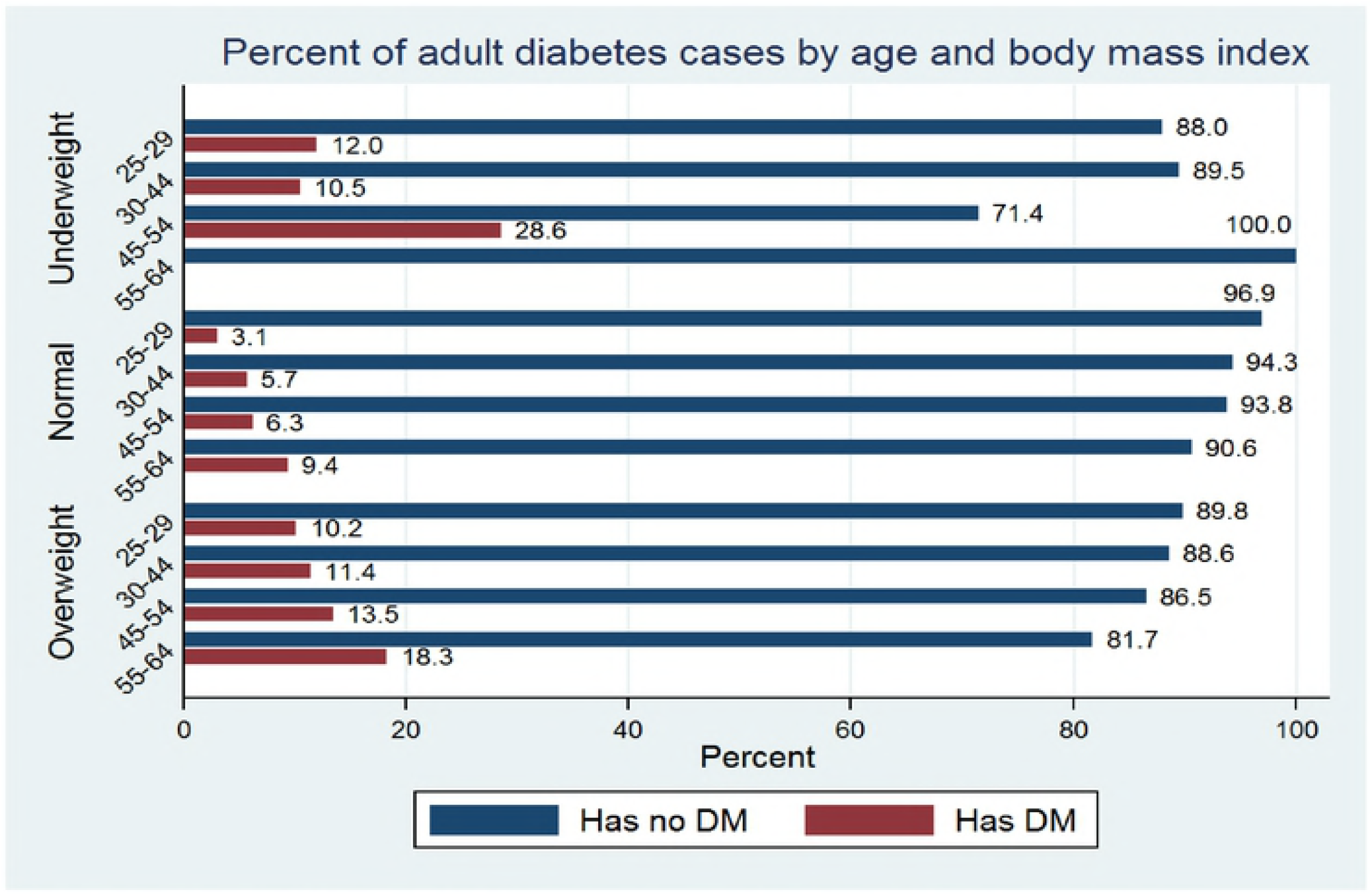
Magnitude of DM by age and BMI among adults age 25-65 in Diredawa City, 2017.

### Factors associated with diabetes mellitus

Three binary logistic regression models were run to identify factors associated with DM considering risk factors function at different levels. At each level variable with p-value less 0.3 was considered for Hierarchical logistic regression model.

**The model I** was adjusted for social structures. The odds of Diabetic Mellitus was 2.4 (AOR 95% CI: 1.1, 5.2) among the age group of 55-64.

**Model II** was adjusted for social positions and behavioral characteristics. The odds of Diabetic Mellitus was 2.3(AOR 95% CI: 1.1, 5.0) among the age group of 55-64.

**Model III** was adjusted for social positions, behavioral characteristics and physical and medical measurement. Consuming two or less serving of vegetables/week, 2.1 (AOR 95% CI: 1.1, 2.9) and having normal BMI, 0.6(AOR 95% CI: 0.3, 0.8) was significantly associated with DM.

**Table 4:** Multi-level correlates of diabetes mellitus among adults in Dire Dawa, Eastern Ethiopia, 2017

| Characteristics |  | DM |  | Model 1<br>AOR(CI) | Model 2<br>AOR(CI) | Model 3<br>AOR(CI) |
| --- | --- | --- | --- | --- | --- | --- |
|  |  | Yes<br>(n=70) | No<br>(n=712) |  |  |  |
| **Sex** | Male | 20(28.6) | 240(33.7) | 0.9(0.4, 1.3) | 0.7(0.4, 1.3) | 0.8(0.4, 1.4) |
|  | Female | 50(71.4) | 472(66.3) | Ref. | Ref. | Ref. |
| **Age** | 25-29 | 13(18.6) | 201(28.2) | Ref. | Ref. | Ref. |
|  | 30-44 | 24(34.3) | 267(37.5) | 1.4(0.7, 2.9) | 1.3(0.6, 2.7) | 1.2(0.5, 2.5) |
|  | 45-54 | 12(17.1) | 95(13.4) | 2.1(0.9, 5.0) | 2.0(0.8, 4.8) | 1.6(0.6, 4.0) |
|  | 55-64 | 21(30.0) | 149(20.9) | **2.4(1.1, 5.2)\*** | **2.3(1.1, 5.0)\*** | 2.0(0.9, 4.4) |
| **Marital status** |  |  |  |  |  |  |
|  | Married | 53(75.7) | 467(65.6) | Ref. | Ref. | Ref. |
|  | Never married /Single | 11(15.7) | 145(20.4) | 0.9(0.4, 1.8) | 0.9(0.4, 1.9) | 0.9(0.4, 1.9) |
|  | Divorced | 3(4.3) | 63(8.8) | 0.4(0.1, 1.2) | 0.4(0.1, 1.2) | 0.4(0.1, 1.3) |
|  | Widowed | 3(4.3) | 37(5.2) | 0.5(0.1, 1.7) | 0.5(0.1, 1.7) | 0.5(0.1, 1.7) |
| **Ever drink alcohol** |  |  |  |  |  |  |
|  | Yes | 23(32.9) | 216(30.3) |  | 1.3(0.7, 2.2) | 1.2(0.7, 2.1) |
|  | No | 47(67.1) | 496(69.7) |  | Ref. | Ref. |
| **Ever smoke tobacco products** |  |  |  |  |  |  |
|  | Yes | 7(10.0) | 60(8.4) |  | 1.4(0.6, 3.4) | 1.4(0.6, 3.5) |
|  | No | 63(90.0) | 652(91.6) |  | Ref. | Ref. |
| **Physical Activity** |  |  |  |  |  |  |
|  | ≤ 600 MET-minutes | 34(48.6) | 290(40.7) |  | 1.3(0.8, 2.2) | 1.3(0.8, 2.1) |
|  | >600 MET-minutes | 36(51.4) | 422(59.3) |  | Ref. | Ref. |
| **Fruit consumption** |  |  |  |  |  |  |
|  | twice or fewer servings/week | 22(31.4) | 264(37.1) |  | 0.6(0.2, 1.1) | 0.8(0.4, 1.4) |
|  | 3-4 servings/week | 15(21.4) | 150(21.1) |  | 0.9(0.5, 1.8) | 0.9(0.4, 1.9) |
|  | 5 or more servings /week | 33(47.2) | 298(41.8) |  | Ref. | Ref. |
| **Vegetable consumption** |  |  |  |  |  |  |
|  | twice or fewer servings/week | 23(32.9) | 229(32.1) |  | 1.8(0.9, 2.8) | **2.1(1.1, 2.9)\*** |
|  | 3-4 servings/week | 18(25.7) | 212(29.8) |  | 0.8(0.4, 1.4) | 0.7(0.4, 1.3) |
|  | 5 or more servings /week | 29(41.4) | 271(38.1) |  | Ref. | Ref. |
| **HTN** |  |  |  |  |  |  |
|  | Yes | 22(34.1) | 148(79.2) |  |  | 1.4(0.8, 2.4) |
|  | No | 48(68.6) | 564(79.2) |  |  | Ref. |
| **Body mass index** |  |  |  |  |  |  |
|  | Underweight | 7(10.0) | 58(8.1) |  |  | 0.9(0.4, 2.2) |
|  | Normal | 24(34.3) | 397(55.8) |  |  | **0.4(0.3, 0.8)\*** |
|  | Overweight | 39(55.7) | 257(36.1) |  |  | Ref. |

Ref. => Reference category
\*= p<0.05, statistically significant

## Discussion

This study found that 8.95% (95% CI: 7.1, 11.2) of adults aged 25-64 years had DM and 3.3% (95% CI: 2.3, 4.8) had never diagnosed for DM. This study considered a multi-level nature of covariates and the analysis revealed that age, vegetable consumption, and BMI were significantly associated with DM.

Our findings on the magnitude of DM among adult people were comparable to sub-national studies conducted in different parts of Ethiopia [9, 12, 18, 29] ranging from 5.1% in Northwest Ethiopia to 11.5% in East Gojam, and 5.9% at national level. The finding was also close to different subnational studies in Africa [11, 17, 20–24, 26], reported a magnitude ranging from 1.4% in Uganda to 19.1% in North Sudan. Our finding, however, is relatively lower compared to the prevalence of DM in a sub-national sample of Asian populations including the India industrial communities, 10.1% [44], Saudi Arabia, 12.1%[25], Iran, 12.3% [19] and, Indonesia, 13% [13]. The difference in magnitude might be due to the rapid rise in the prevalence of DM[6], in addition to the geographical difference and an associated lifestyle [9], change in sedentary lifestyle and urbanization[45] and, the low level of physical activity associated with urban lifestyle in the current study [46].

In this study, 3.3% of the total participant or 36.9% of identified diabetic cases were undiagnosed for DM prior to the survey. The finding was consistent with sub national studies of 32.7% [12] and study conducted across Africa, ranging from 31.3%[22] in North Sudan to 34.5% in Zambia [11]. The finding is also comparable with study report of sub national studies in Asia including Indian industrial community, 38.4%[44] and central Vietnam, 44.6%[47]. Thus, efforts have to be made, particularly by the individuals involved in health practice, to early detect the disease and thereby initiate a suitable therapeutic service, before complications, arise. Contrary to the above report, the magnitude of undiagnosed DM was relatively lower compared to the study report of developing countries which reported 65.2% magnitude of undiagnosed DM [48] and global estimate of 50%[3]. The lower magnitude of undiagnosed DM in the current study could be relatively due to better health service coverage and health service utilization in the study area[49].

The recent study also found that the prevalence of DM was higher among those of an older age. This is in line with the previous studies conducted around the globe [11–15, 18, 19, 21–25, 28, 44]. This is likely due to people’s tendency to exercise less, lose muscle mass and gain weight as people get age[50]. In addition, pancreatic β-cell mass in adults exists in a dynamic state such that the cells can undergo compensatory changes to maintain euglycemia. However, as an individual gets older the capacity of β-cells regeneration is reduced. In another way, aging has direct effects on β-cell proliferation and function that contributes indirectly to impaired insulin sensitivity mediated by lifestyle and comorbidity-related risk factors[51]. Therefore, with an increase in age, associated DM prevention measures might include, healthy diet, regular physical activity, maintaining normal body weight and avoiding alcohol and tobacco use[5].

Similar to the finding of previous studies [11, 13–17, 19, 21–24, 27, 29], the occurrence of DM is higher among overweight and obese adults. This might be due to the fact that overweight and obese individuals’, adipose tissue releases increased amounts of non-esterified fatty acids, glycerol, hormones, pro-inflammatory cytokines and other factors that are involved in the development of insulin resistance. Insulin resistance is accompanied by dysfunction of pancreatic islet β-cells the cells that release insulin to control blood glucose levels. Abnormalities in β-cell function are therefore critical in defining the risk and development of DM [52]. Therefore, routine public education campaigns should be conducted to create awareness about lifestyle modification and the necessity of weight reduction.

Consistent with previous research report [30–33, 53–55] the current study reported, frequent use of vegetables reduce the risk of DM. Vegetables are rich sources of antioxidant compounds such as carotenoids, vitamin C, vitamin E and flavonoids, and of fiber and may have a protective effect against the development of diabetes by relieving oxidative stress that interferes with the glucose uptake by cells[56]. It has also been hypothesized that soluble fibers from vegetables inhibit the postprandial glucose load by delaying the absorption of carbohydrates and may thus inhibit diabetes development[57]. However, we did not measure the type of vegetables that participants consumed and data collection was based on self-reporting which might have biased our measurement, and subsequently the estimate.

### The strength

A key strength of the study was that that the chronic nature of diabetes mellitus has been considered along with causal pathways, even though, it is not specific to DM, we have tried to identify the covariates linked with DM at different hierarchical levels (distal to proximal) instead of simply including as many covariates as possible to search for the ones that are significantly associated. In addition, the study was conducted at population level on a representative sample and blood sugar was measured instead of relying on self-report so that previously undiagnosed raised blood sugar cases were identified.

### Limitations

While sharing the methodological limitation of cross-sectional studies with others, the effects of income or wealth, family history, and other eating behaviors other than fruits and vegetables are not assessed. Unintentionally, the majority of study participants were female and this may bias the overall estimation. The study also did not consider acute and chronic illnesses which may affect the blood sugar level.

### Conclusion

The overall magnitude of diabetes mellitus among adults in Dire Dawa was high and the magnitude of undiagnosed DM was a great concern. Age, BMI, and frequency of vegetable consumptions are significant risk factors for the DM. Therefore, awareness creation activities targeting adults should be devised and institutionalized in the health system. Campaign and community mobilization to sensitize the community about early screening and preventive packages targeting modifiable risk factors should be employed to reduce the burden of diabetes mellitus.

## Abbreviations

AOR: adjusted odd ratio
BMI: body mass index
BP: blood pressure
CI: confidence interval
DM: diabetes mellitus
FBS: fasting blood sugar
HU: Haramaya University
IDA: International Diabetes Federation
IHRERC: Institutional Health Research Ethics Review Committee
LMIC: low- and middle-income countries
MICE: multiple imputations by chained equations
NCD: non-communicable diseases
RBS: random blood sugar
SD: standard deviation
USD: United state dollar
WHO: World health organization

## Acknowledgment

We would like to thank Haramaya University for providing us with financial support. We would like to extend our gratitude to Dire Dawa City Administration Health Bureau and to the community. We would also like to thank Ms. Rasha Darghawth for her diligent language edition.

